# Early life sleep disruption has long lasting, sex specific effects on later development of sleep in prairie voles

**DOI:** 10.1101/2022.10.18.512732

**Authors:** Carolyn E. Jones-Tinsley, Randall J. Olson, Miranda Mader, Peyton T. Wickham, Katelyn Gutowsky, Claire Wong, Sung Sik Chu, Noah E. Milman, Hung Cao, Miranda M. Lim

## Abstract

In mammals, sleep duration is highest in the early postnatal period of life and is critical for shaping neural circuits that control the development of complex behaviors. The prairie vole is a wild, highly social rodent that serves as a unique model for the study of complex, species-typical social behaviors. Previous work in our laboratory has found that early life sleep disruption (ELSD) in prairie voles during a sensitive window of postnatal development leads to long lasting changes in social and cognitive behaviors as well as structural changes in excitatory and inhibitory neural circuits in the brain. However, it is currently unknown how later sleep is impacted by ELSD, both shortly after ELSD and over the long term. Therefore, the aim of this study was to describe the effects of ELSD on later life sleep, compared to sleep in normally developing prairie voles. First, we conducted tethered electroencephalogram/electromyogram (EEG/EMG) recordings in juvenile prairie voles undergoing ELSD, compared to Control conditions. Second, we conducted 24 hours of home cage tethered EEG/EMG recordings in either adolescent or adult male and female prairie voles that had previously undergone ELSD or Control conditions as juveniles. We found that, as adults, male ELSD prairie voles showed persistently lower REM sleep duration and female ELSD prairie voles showed persistently higher NREM sleep duration compared to Controls, but no other sleep parameters differed. We concluded that 1) persistent effects of ELSD on sleep into adulthood may contribute to the social and cognitive deficits observed in adult voles, and 2) sleep disruption early in life can influence later sleep patterns in adulthood.

## INTRODUCTION

In mammals, sleep duration, and in particular, rapid eye movement (REM), is highest in the early postnatal period of life. This period of increased REM may be necessary to shape neural circuits that control the development of complex behaviors, such as species-typical social behaviors. Prairie voles (*Microtus ochrogaster*) are an altricial rodent species commonly used to study social behavior. In both the wild and the laboratory, prairie voles are believed to form long-lasting pair bonds with opposite sex mates, making them an ideal model species to study the underlying neural circuitry of social bonding [1, 2], as well as the various genetic and environmental factors that affect formation and maintenance of social bonds [3, 4]

We have previously shown that early life sleep disruption (ELSD) during a sensitive window in prairie vole development (postnatal week 3) results in long lasting changes in both social and cognitive behavior as adults [5, 6]. Our method of ELSD during postnatal day P14 to P21 using automated, gentle cage agitation at timed intervals produced selective reduction in rapid eye movement (REM) sleep, as well as fragmentation in non-REM (NREM) sleep, while allowing pups to remain otherwise undisturbed in their home cages without significant alterations stress hormones or quantity of parental care received [6].

We found that ELSD resulted in long-term changes in social and cognitive behaviors in prairie voles as adults, including reduced affiliative huddling during the partner preference test (e.g., widely regarded as a laboratory proxy for prosocial aspects of the pair bond), as well as impaired fear extinction (a proxy for cognitive flexibility) [5, 6]. At the neuronal level, and consistent with the social and cognitive behavioral deficits above, we reported that ELSD altered markers of both excitation and inhibition within the cortex, including increased immunoreactivity of inhibitory parvalbumin positive interneurons within the primary somatosensory cortex [6], and increased dendritic spine density in layer 2/3 of prefrontal cortex [5].

However, it is currently unknown how ELSD alters sleep during postnatal week 3 and whether ELSD results in long-term changes in sleep in prairie voles. Understanding the long-term effects of ELSD on sleep later in life would provide insight into the factors that shape the development and neural control of healthy sleep. Furthermore, characterizing the normal ontogeny of sleep in developing prairie voles would provide insight into the underlying biology of sleep in this naturally occurring, highly social species, and could also potentially enhance generalization and translation of results to human sleep and behavior. In order to address these scientific gaps, the aim of this study was to examine the effects of ELSD in prairie voles on their sleep patterns later in life.

## METHODS

Using skull-based EEG combined with nuchal EMG, we collected objective sleep measures analogous to the human polysomnography at 3 time points in prairie vole development: 1) during early life sleep disruption (postnatal week 3), 2) early adolescence (7 days after Early Life Sleep Disruption (ELSD)), and 3) young adulthood (7 weeks after ELSD). All procedures were approved by the Institutional Animal Care and Use Committee at the Portland VA Medical Center and were conducted in accordance with the National Institutes of Health Guide for the Care and Use of Laboratory Animals.

### Experimental Design

Data was collected across two experiments. In Experiment 1, juvenile prairie voles were implanted with EEG/EMG electrodes and sleep was recorded for 6 consecutive days in the midst of either ELSD (n=2 female and n=3 male) or Control (n=3 female and n=2 male) conditions. This experiment was conducted in order to describe the full sleep phenotype produced by ELSD, including any potential habituation to the ELSD paradigm, in young voles. In Experiment 2, a separate cohort of ELSD (n=6 female and n=8 male) or Control (n=5 female and n=5 male) prairie voles were implanted with EEG/EMG electrodes and sleep was recorded for 24 hours during early adolescence (or approximately 7 Days after the completion of ELSD or Control sleep manipulation). In Experiment 3, ELSD (n=9 female and n=10 male) or Control (n=9 female and n=10 male) prairie voles were implanted with EEG/EMG electrodes and sleep was recorded for 24 hours during adulthood (approximately 10-15 weeks after the completion of ELSD or Control sleep manipulation).

### Subjects

Subjects were male and female prairie voles bred in house at the VA Portland Health Care System and reared by both parents. Subjects were housed in clear polycarbonate cages (27 cm × 27 cm × 13 cm) and housed in temperature and humidity controlled rooms on an automated 14:10 light:dark cycle (lights on at 0700h). Animals had *ad libitum* access to a mixed diet of rabbit chow (LabDiet Hi-Fiber Rabbit), corn (Nutrena Cleaned Grains), and oats (Grainland Select Grains) and water (water bottles and/or hydrogel) throughout the entirety of both experiments. Cotton nestlets and a wooden block or stick for chewing enrichment were added to each cage and replaced weekly with cage change. No litter contributed more than n=2 animals of the same sex to any given experiment. The prairie vole colony originated from Emory University derived from field caught prairie voles in Illinois. Colony diversity was maintained through generous bi-annual donations from researchers across the United States, including Dr. Lisa McGraw at North Carolina State University in 2014, Dr. Karen Bales at UC Davis in 2015, Dr. Zoe Donaldson at CU Boulder in 2017, and Dr. Zuoxin Wang at Florida State University in 2019. Breeder pairs were checked each morning at lights on for the presence of pups and the day of pup discovery was designated as postnatal day (P)1. Voles were weaned at P20 and socially housed with same sex littermates (2-4/cage) until testing (P20, ∼P29, ∼P100).

In Experiment 1, two voles were removed from analysis because they did not produce a full 6 days of usable sleep data due to equipment malfunction.

### Early Life Sleep Disruption

To disrupt sleep early in life (ELSD group), home cages containing prairie vole litters with both parents present were placed on a standard laboratory orbital shaker connected to a timer programmed to turn on every 110 seconds for 10 seconds, thus providing gentle agitation to the entire cage (110 rotations per minute). With this method, pups remain otherwise undisturbed, parental care is not disrupted, and markers of stress are not increased [6]. Cage card holders were locked into place to limit auditory disruption during cage agitation. Water bottles were removed and hydrogel was provided as an alternative to prevent spillage during shaking. Cages containing control animals were housed in the same room as the shakers and supplemented with hydrogel but cages were not physically disturbed. In Experiment 1, sleep was recorded during ELSD, as such, young voles were housed individually on the orbital shaker along with support fluids and softened food.

### EEG/EMG electrode implantation

Custom made electrodes were built for each animal prior to implantation. Electrodes consisted of an 8-pin female header socket (MillMax) with 4 silver EEG leads (A-M Systems, Inc) soldered to the top row of pins and 3 insulated silver wire EMG leads soldered to the bottom row. To implant electrodes, voles were anesthetized with isoflurane (3-5% induction, 1-3% maintenance) and affixed in a stereotaxic frame (Kopf) customized for prairie voles. During surgery, the skull was exposed and 4 small holes drilled (2 frontal and 2 parietal) for the insertion of stainless-steel screws soldered to the EEG wires. EMG wires were partially stripped of their insulation before being routed through the dorsal neck muscles. The entire electrode was secured to the skull using both dental cement and a small amount of cyanoacrylate glue. The animal received subdermal fluids and carprofen (5 mg/kg, i.p.) before being placed into a clean home cage on top of a heating pad until alert and ambulatory. Support heat, softened food, hydrogel, and extra nestlets were provided during recovery.

### Sleep Recordings

Animals were connected to custom built lightweight recording cables (Cooner wire) via male header sockets and given enough slack to move freely. Data was collected from a Grass amplifier via AcqKnowledge software (BIOPAC). In Experiment 1, sleep signals were recorded for 6 consecutive days starting on P20. In Experiments 2 and 3, animals were recorded for 3 consecutive days starting on approximately either P29 (Experiment 2) or P100 (Experiment 3) with the middle 24 hours used for analysis.

### Sleep Scoring

EEG/EMG signals were converted to European Data Format before being scored offline for NREM and REM sleep and Wake in 4-second epochs across the full 24-hour period (SleepSign, Kissei Comtec). EEG channels were filtered with a high-pass filter at 0.75 Hz and EMG channels with a band-pass filter from 20-50 Hz. Wake was defined as high amplitude mixed frequency EEG paired with high amplitude EMG. NREM was defined by high amplitude, low frequency EEG paired with low amplitude EMG. REM was defined by low amplitude, high frequency EEG paired with low amplitude EMG with occasional muscle twitches. EDF files were scored per the above criteria automatically (SleepSign) and then manually verified and corrected by trained experimenters. Brief arousals were defined as <= 3 epochs (12 sec) of Wake immediately following >= 3 epochs (12 sec) of sleep (NREM+REM). Wake bouts were defined as >=4 epochs of Wake immediately following >=3 epochs of sleep (NREM+REM) (Li et al 2014). Two measures of sleep fragmentation were used following the methods of an identical method of sleep disruption on adult mice (Li et al 2014): 1) Average bout length for NREM and REM sleep and 2) arousal index, which was calculated by combining the total frequency of brief arousals and wake bouts per hour of sleep (NREM+REM) over the full 24 hours of recording for each day.

### Statistics

Assumptions were met for parametric testing and group differences were analyzed with ANOVA (between group factors: Sex, Sleep Group). When the same animal was tested at multiple time points, repeated measures ANOVA were used (within group factor: day or age). If sphericity tests were violated (Mauchly’s), F values were Greenhouse-Geisser corrected. In Experiment 1, we were not powered to detect sex differences and males and females were analyzed together. Significant interactions were followed up with independent sample t-tests (two-tailed) with alpha values set at 0.05.

## RESULTS

### Experiment 1 – 6 days of shaker sleep disruption

#### Young prairie voles acclimate to sleep fragmentation during ELSD

Sleep fragmentation was quantified using a combination of Arousal Index (# Brief arousals + # Wake Bouts/Total Sleep Time) and average NREM and REM bout length for each 24-hour period.

##### Arousal Index

Repeated measures ANOVA (between group factor = sleep group; within subjects factor = recording day) indicated a significant day by sleep group interaction *F*(5,40)=4.022, *p*=0.005.Follow-up independent sample t-tests revealed that arousal index was only increased during ELSD on the first full day that animals were housed on the shaker (*t*(8)=6.202, *p*<0.001) with no significant differences detected on other shaker days (all *p* values >0.15) (**Fig 2A**).

**Figure 1:**
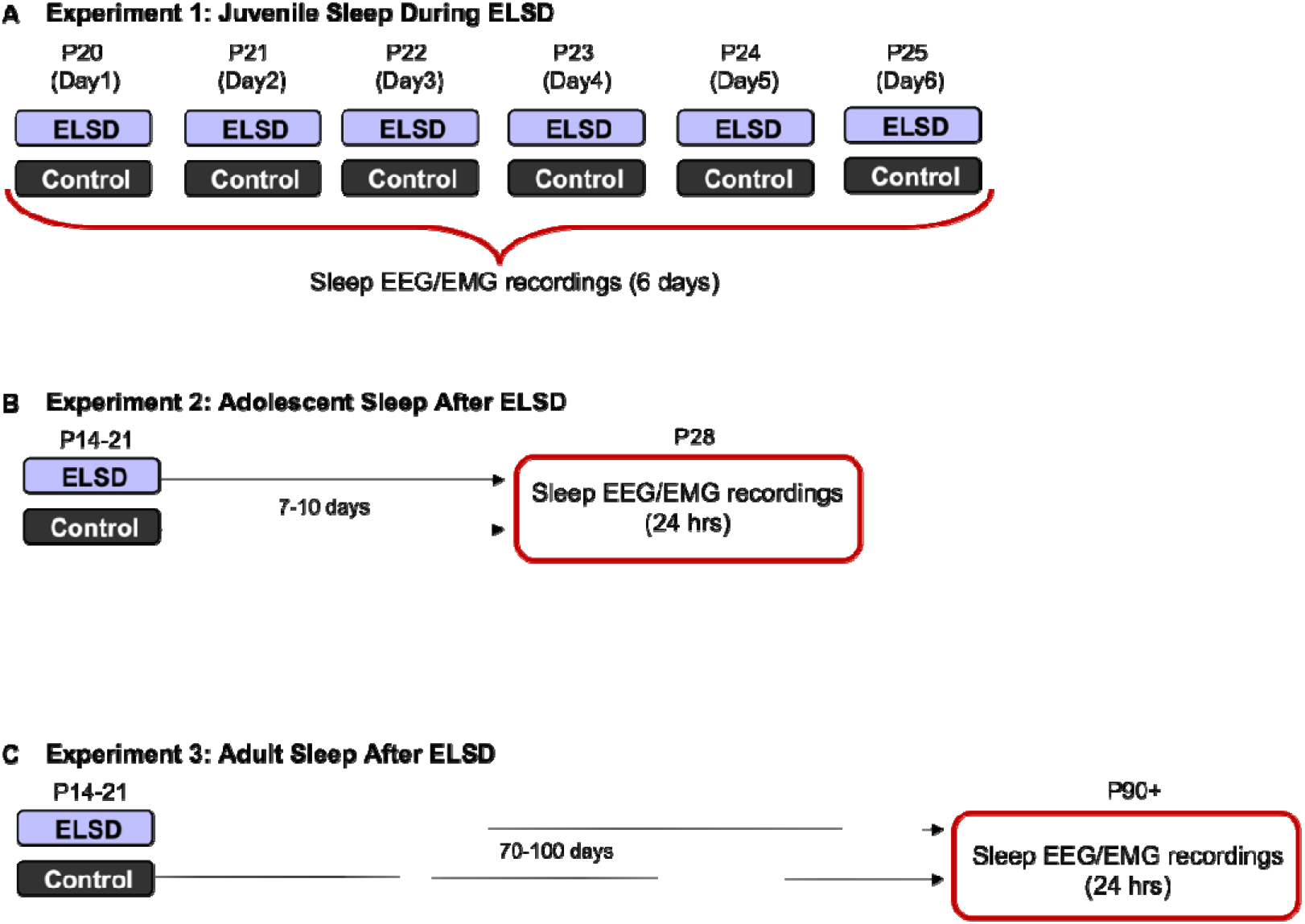
Experimental design. Three experiments were conducted to describe sleep in prairie voles during and after ELSD using EEG and EMG recordings over 24-hour bins during different developmental windows. A) In experiment 1, sleep was recorded during ELSD or Control conditions for 6 consecutive days. B) In experiment 2, sleep was recorded after ELSD or Control conditions during early adolescence. C) In experiment 3, sleep was recorded after ELSD or Control conditions during adulthood.

**Figure 2:**
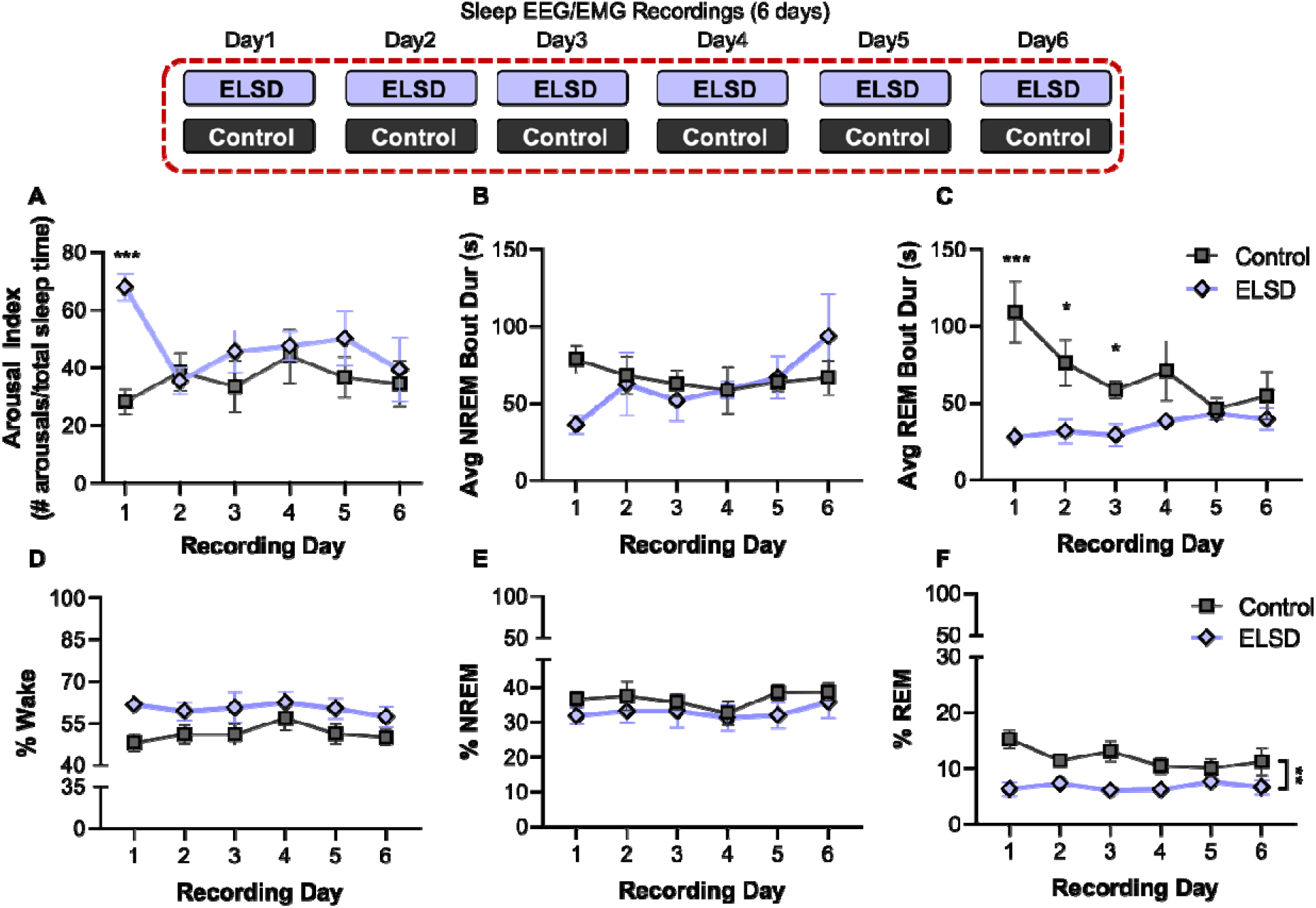
Experiment 1: ELSD in juvenile voles acutely fragments REM and NREM sleep and chronically reduces REM sleep duration. A) Arousal Index (# arousals/total sleep time) was increased only on the first day of ELSD. B) Average length of NREM sleep bout each day of recording was not significantly reduced. C) Average length of REM sleep bout was decreased the first 3 days of ELSD. D) Time spent in Wake across the entire 6 days of ELSD was not significantly increased. E) No change in NREM sleep amounts. F) REM sleep amount were reduced across the entire 6 days of ELSD. N=5/group. Symbol represents mean, error bars +/-SEM. *p<0.05; **p<0.01; ***p<0.001 REM = rapid eye movement; NREM = non-rapid eye movement; ELSD = early life sleep disruption

##### NREM Average Bout Length

There were no significant day by sleep group interactions on NREM average bout length (repeated measures ANOVA, Greenhouse-Geisser corrected: F(2.3,18.3)=2.808, p=0.167) and no significant main effects (between group effect of sleep group: F(1,8)=0.135, p=0.723) (**Fig 2B**).

##### REM Average Bout Length

There was a significant day by sleep group interaction on REM average bout length (repeated measures ANOVA, Greenhouse-Geisser corrected: F(1.84,14.72)=4.37, p=0.035) as well as a main effect of group (between subjects factor: F(1,8)=9.828, p=0.014). Follow up t-tests revealed that the average REM bout length was decreased on the shaker for the each of the first 3 days (day 1: t(8)=3.854, p=0.005; day 2: t(8)=2.622, p=0.031; day 3: t(8)=3.237, p=0.012; day 4: t(8)=1.658, p=0.136; day 5: t(8)=0.402, p=0.698; day 6: t(8)=0.886, p=0.401) (**Fig 2C**).

#### REM sleep duration is reduced for entire ELSD period

Duration in each vigilance state was calculated as a percentage by dividing the total time spent in each state (Wake, NREM, REM) by the total recording time for each day.

For all 3 vigilance states there were no significant within subjects interactions over the 6 days of ELSD (all p values >0.728). There was a trend towards increased time in Wake in animals on the shaker that did not reach significance (repeated measures ANOVA, between subjects effect of sleep group: F(1,8)=5.055, p=0.055) (**Fig 2D**). There was not a significant effect of housing on the orbital shaker on NREM sleep duration (between subjects effect of sleep group: F(1,8)=1.185, p=0.308) (**Fig 2E**). There was a main effect of sleep group on REM sleep duration with voles housed on the shaker during ELSD spending significantly less time in REM sleep compared to controls across the 6 days of recording (between subjects effect of sleep group: F(1,8)=19.212, p=0.002) (**Fig 2F; see Tables S1 and S2** for mean REM sleep duration by sex).

### Experiment 2: ELSD does not affect adolescent sleep

#### Sleep fragmentation

There were no long term effects of Early Life Sleep Disruption on any of the sleep fragmentation metrics quantified here at adolescence (main effect of sleep group-arousal index: F(1,20)=1.29, p=0.270 (**Fig3a**); NREM Avg Bout: F(1,20) = 0.435, p=0.517 (**Fig 3b**); REM avg bout F(1,20)=0.021, p=0.886 (**Fig 3c**)). There were also no significant effects of sex on sleep fragmentation in prairie voles in adolescence (arousal index, main effect of sex: F(1,20)=0.002, p=0.961; NREM avg bout F(1,20)=0.042, p=0.839; REM avg bout F(1,20)=0.013, p=0.911) and none of the group x sex interaction terms were significant (all p values >0.709)

**Figure 3:**
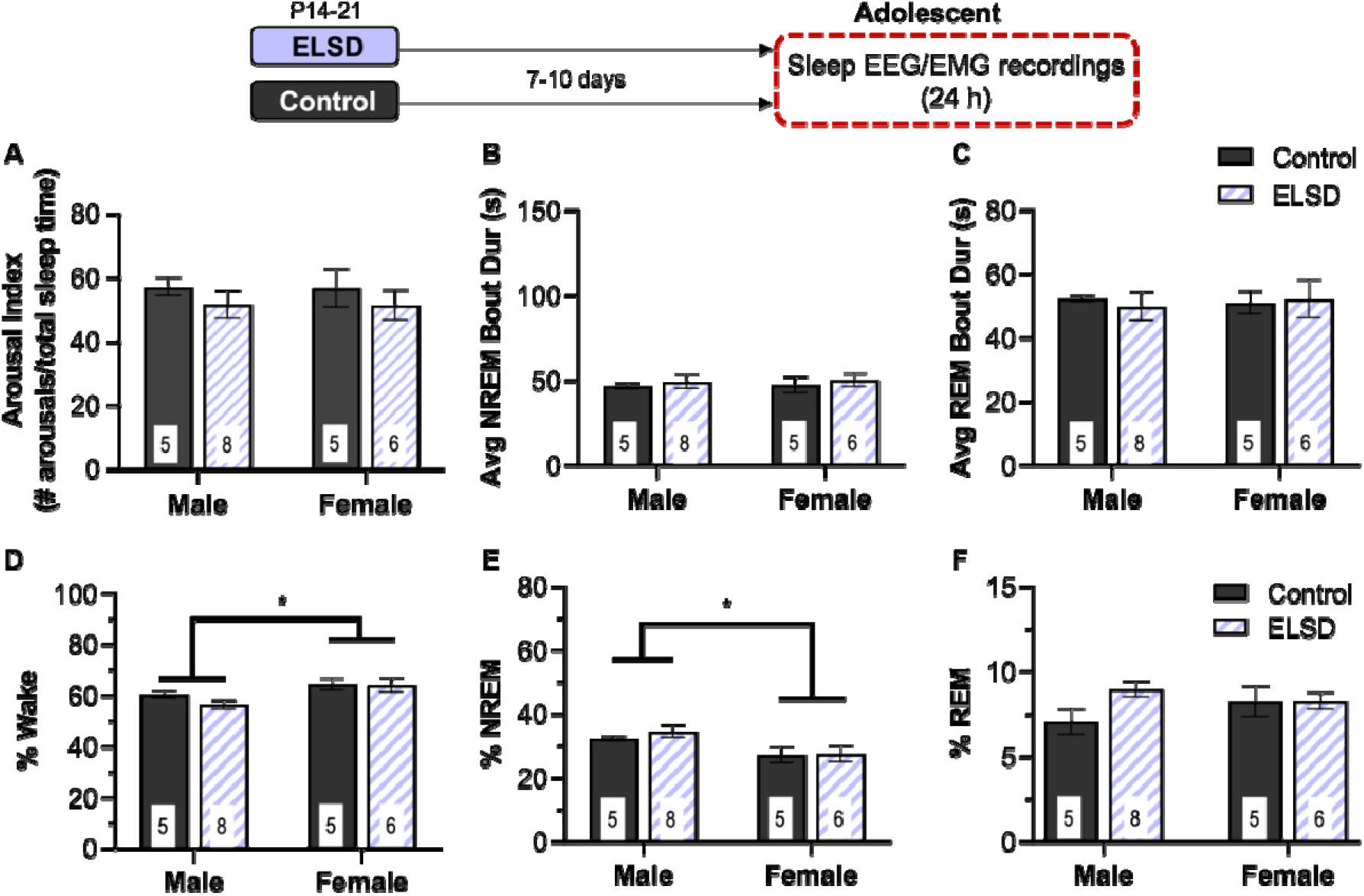
Experiment 2: Adolescent sleep was not altered after ELSD. There were no long term effects of ELSD on measures of sleep fragmentation during early adolescence including arousal index (# arousals/total sleep time) (A), NREM average bout length (B), or REM average bout length (C). Adolescent females spent more time in wake than males (D) and less time in NREM (E) but these were not affected by the ELSD procedure. F) there were no significant changes in REM sleep time during adolescents after ELSD compared to Controls. Bar height is mean, number in bars is sample size, error bars +/-SEM. *P<.0.05

#### Wake Duration

ELSD did not have a significant effect on time spent in wake during adolescence (main effect of sleep group: F(1,20)=0.952, p=0.341) (**Fig 3d**). Adolescent females spent more time in wake than adolescent males (main effect of sex: F(1,20)=6.623, p=0.018) and the interaction was not significant (F(1,20)=0.689, p=0.416).

#### NREM Sleep Duration

Adolescent female prairie voles spent significantly less time in NREM sleep than males (main effect of sex: F(1,20)=7.576, p=0.012)(**Fig 3e**). There were no significant effects of early life sleep group on NREM sleep duration during adolescence (main effect of sleep group: F(1,20)=0.385, p=0.542; sleep group x sex interaction: F(1,20)=0.232, p=0.635).

#### REM Sleep Duration

The effects of ELSD on REM sleep duration in adolescent prairie voles was not significant (main effect of sex: F(1,20)=0.162, p=0.692; main effect of sleep group: F(1,20)=2.405, p=0.137; sleep group x sex interaction: F(1,20)=2.266, p=0.148)(**Fig 3f**).

### Experiment 3: ELSD reduces REM sleep duration in adult males

#### Sleep fragmentation

There were also no long term effects of Early Life Sleep Disruption on any of the sleep fragmentation metrics quantified here at adulthood (main effect of sleep group-arousal index: F(1,34)=0.808, p=0.375 (**Fig 4a**); NREM Avg Bout: F(1,34)=0.047, p=0.829 (**Fig 4b**); REM Avg Bout: F(1,34)=2.298, p=0.139) (**Fig 4c**). However, in contrast to adolescence, in adulthood, females had more consolidated NREM sleep than males as evidenced by reduced arousal index (main effect of sex: F(1,34)=6.173, p=0.018) and increased average NREM bout length (F(1,34)=4.565, p=0.04) but not average REM bout length (main effect of sex: F(1,34)=1.243, p=0.273 (**Fig 4c**)). Consistent with adolescence, there were no significant sex by early life sleep interactions in adulthood (all p values >0.219).

**Figure 4:**
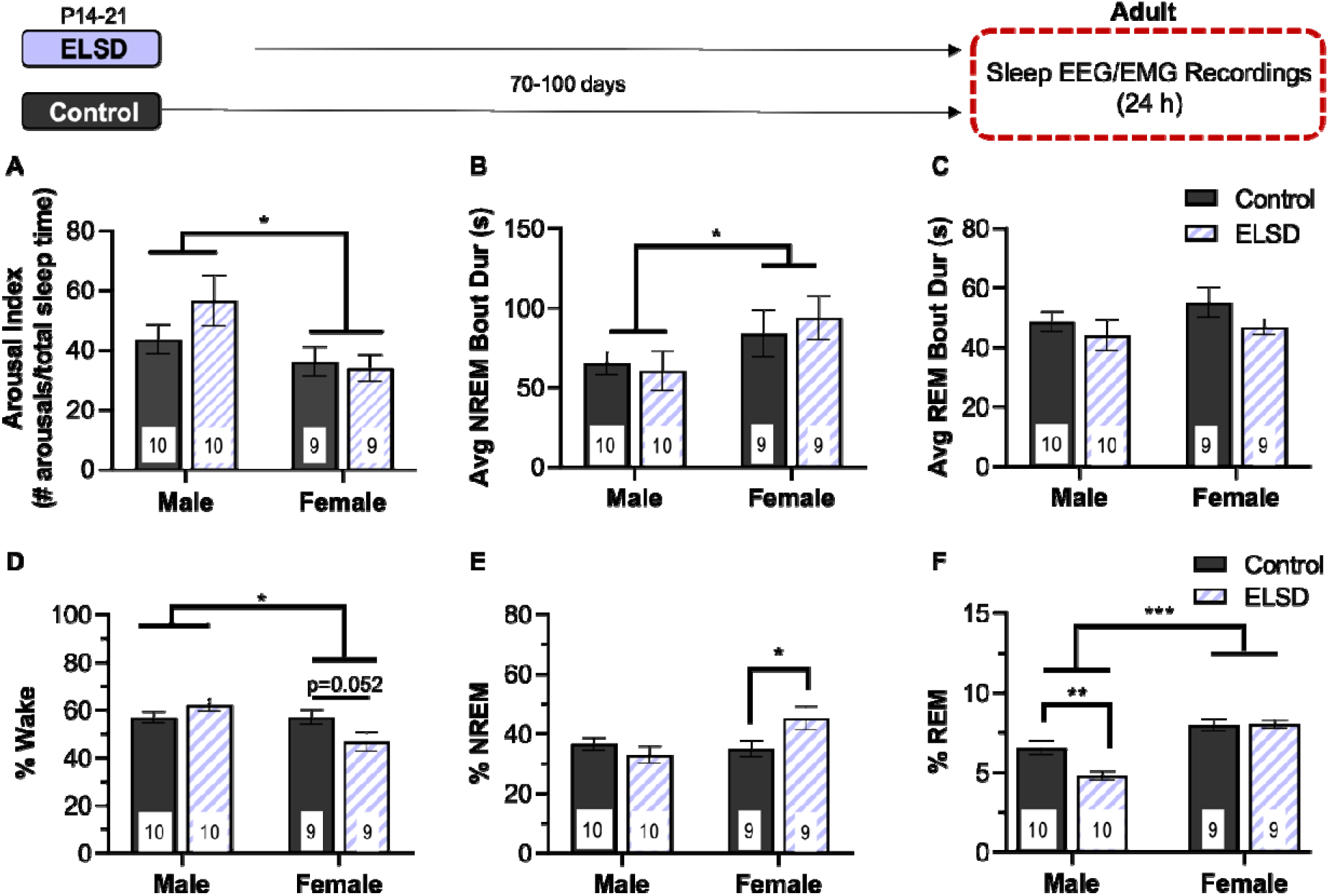
Experiment 3: Persistent effects of ELSD on sleep duration are sex specific and age dependent. A-C) There were no significant effects of ELSD on adult sleep fragmentation metrics including arousal index (A), average NREM bout duration (B), or average REM bout duration (C). D) Adult females spent less time in wake than males. E) ELSD females spent more time in NREM sleep than Control females during adulthood and F) male ELSD voles spent less time in REM sleep than age and sex matched controls. Adult females spent more time in REM than adult males and have more consolidated NREM sleep (B). Note y-axis limit is different on each panel. *p<0.05; **p<0.01; ***p<0.001; number at bottom of bar is sample size for each group, bar height is mean, error bars +/-SEM

#### Wake Duration

In adulthood, there was not a significant main effect of sleep group: F(1,34)=0.658, p=0.423 and adult females spent less time in wake than adult males (main effect of sex: F(1,34)=6.448, p=0.016). There was a significant sleep group by sex interaction in adult prairie voles (F(1,34)=6.710, p=0.014) but follow up t-tests did not reach significance (males t(18)=1.461, p=0.161; females t(16)=2.101, p=0.052) (**Fig 4d**).

#### NREM Sleep Duration

In contrast to sex effects on NREM sleep observed in adolescence, there were no main effects of sex on NREM sleep duration in adults (main effect of sex: F(1,34)=2.191, p=0.148). There was a significant sleep group x sex interaction in adults (F(1,34)=6.103, p=0.019) with adult females spending more time in NREM sleep after ELSD than Controls (t(16)=2.226, p=0.041) (**Fig 4e**). We then looked at NREM sleep duration by hour and found a significant hour by sex interaction (repeated measures ANOVA, f(23,759)=1.804, p=0.012) but no within subjects effects of ELSD on REM sleep (all p values >0.531) (**Fig S1**).

#### REM Sleep Duration

Adult females spent significantly more time in REM sleep than adult males (main effect of sex: F(1,34)=32.281, p<0.001) and REM sleep was reduced after ELSD in males only (sleep group x sex interaction: F(1,34)=9.734, p=0.004; t(18)=3.224, p=0.005) (**Fig 4f**). We then looked at REM sleep duration by hour and found a significant hour by sex interaction (repeated measures ANOVA, f(23,759)=2.019, p=0.003) but no within subjects effects of ELSD on REM sleep (all p values >0.325) (**Fig S1**).

## DISCUSSION

In these experiments, prairie vole pups underwent ELSD or Control conditions for one continuous week in postnatal development (postnatal week 3). We conducted sleep EEG/EMG recordings at three separate developmental timepoints in order to describe the effects of ELSD on sleep behavior later in life. We found that during the ELSD period itself, REM sleep quality and quantity are preferentially disrupted. Furthermore, we found that ELSD generates long-lasting effects on REM sleep development in male prairie voles. These findings may be relevant to human neurodevelopmental disorders such as autism spectrum disorder (ASD) that feature impairments in both sleep and social behavior that persist into adulthood.

### Ontogeny of sleep in prairie voles mirrors other altricial rodent species with notable sex differences

To our knowledge, this study is the first description of prairie vole REM and NREM sleep across development. Consistent with other studies of mammalian sleep ontogeny, including other altricial rodent species [7–9], as well as humans [10], as young prairie voles grew older, they spent less time in REM sleep and NREM sleep became more consolidated. Interestingly, we observed several notable sex differences in sleep in prairie voles at both adolescent and adult time points. For example, female adolescent voles spent significantly more time in Wake and less time in NREM sleep than male adolescent voles. Although these sex differences in themselves deserve further exploration, they were not impacted by ELSD *per se*.

Furthermore, we found that adult female voles spent more time in REM sleep than adult male voles and their timing of both REM and NREM sleep throughout the day differed from males. This finding was surprising, and is in contrast to other published literature on rats and mice reporting either decreased REM sleep amounts in females compared to males [11, 12] or no sex differences in REM sleep amounts [13]. REM sleep amounts are influenced by the estrous cycle in female rats [12, 14–17] but the effects of estrus on female mice appears to be minimal and strain specific [18]. Unlike mice and rats, prairie voles are male induced ovulators, and are assumed to be in a persistent state of diestrus if not exposed to males. Also, adult female voles showed less arousals and longer NREM bouts compared to adult male voles, suggesting sex differences in NREM sleep consolidation.

### During the ELSD paradigm itself, sleep fragmentation is only transiently increased, but total REM time is persistently decreased at the expense of Wake

Previous work in our lab recorded sleep on the shaker for 1 day in young prairie voles and reported that ELSD reduces REM sleep time and fragments NREM sleep compared to 1 day of recordings in that same animal off the shaker [6]. Here, we expanded these findings in two ways: 1) we recorded from younger animals more closely matched to the experimental age of our other ELSD experiments and 2) we recorded EEG/EMG for 7 consecutive days (6 full 24-hour periods for analysis) from age-matched animals during either ELSD or Control conditions.

Consistent with our previous work, we found that juvenile prairie voles housed on an orbital shaker experienced significantly reduced REM sleep duration compared to Control voles [6]. During ELSD in prairie voles, REM average sleep bout lengths were shorter than controls, but only transiently so. Furthermore, using an arousal index [19] to quantify arousals from sleep, we found that the orbital shaker only increased arousals for the first day of sleep disruption in young prairie voles. This is in contrast to results reported by Li et al 2014 in adult mice sleep disrupted for 4 continuous weeks. Our method of sleep disruption is identical to the method described by Li et al 2014 in adult mice, however, Li et al found chronic sleep fragmentation, resulting in shorter sleep bouts and increased arousal index sustained across the full period of sleep disruption. Here we report that juvenile prairie voles acclimate to the sleep fragmentation effect of ELSD but do not acclimate to the REM sleep reduction. This may be specific to prairie voles as a species, or it may be specific to this period in early life.

### ELSD cause both age- and sex-specific changes in sleep later in life

We examined prairie voles at two time points after ELSD or Control conditions, and found that ELSD caused long-lasting effects on REM sleep later in life – but only in males. ELSD did not affect sleep patterns in adolescent animals. However, ELSD caused decreased REM sleep in adult males, compared to age- and sex-matched controls. Effects of ELSD on REM sleep females were not significantly different between groups, but there were significant increases in NREM sleep time compared to age- and sex-matched Controls. Notably, ELSD effects on REM sleep were absent in females.

The timepoint chosen for adult sleep phenotyping in these studies mirrors the timing of our previous work describing social and cognitive deficits in adult males after ELSD. These results provide further support of the importance of early life REM sleep on the development of, not only social and cognitive behaviors, but also sleep behaviors. Despite transient effects of sleep fragmentation during the first day of ELSD, this effect resolved over the next 6 days of the ELSD paradigm while total REM time remained persistently decreased over the entirety of the weeklong ESLD protocol.

The exact role that ELSD plays in shaping the underlying neural circuitry relevant to both sleep and social behavior is still unknown. It is still unknown whether the combination of ontogenetic REM sleep changes observed in male prairie voles that underwent ELSD leads to long term REM sleep reduction in adulthood and subsequent impairments in social bonding behavior or if long lasting REM sleep disturbance exist after ELSD independent of social and cognitive impairments.

### Possible relevance of early life sleep disruption to human ASD

There is a large and growing body of research describing sleep problems in patients diagnosed with ASD. The majority of this work describes difficulties with initiating and/or maintaining sleep [20–22] as well as decreased time in REM sleep [23, 24]. This early life sleep phenotype in children with ASD mirrors the sleep disturbance experimentally created with the ELSD paradigm in the current study.

In patients with ASD, sleep problems persist throughout development and into adulthood [25, 26] and may increase as young children grow older [27, 28]. Research examining sleep in older children and adolescents with ASD suggests that in addition to problems maintaining sleep at night, this age group also reports a high frequency of daytime sleepiness and oversleeping [29, 30], which may be consistent with our findings indicating that adolescent voles previously subjected to ELSD do not show reduced total REM sleep times compared to age and sex matched controls. Adults with ASD reveal problems falling asleep (longer sleep latency) and frequent night awakenings as well as reduced time in NREM sleep and reduced rapid eye movements during REM sleep. [31]. While our ELSD paradigm did not result in long term reductions in NREM sleep, ELSD males (but not females) spent less time in REM sleep than their age and sex matched counterparts. Finally, other research has pointed to adolescence being a unique time that does not necessarily show linearity in other behavioral trajectories (reviewed in [32]).

Research related to sex differences in sleep and ASD is mixed. While some studies report increased sleep disturbances in females with ASD throughout development [33, 34] others suggest that male sex contributes to the severity of sleep disruption or the association between sleep and ASD symptomology [27, 35, 36]. Our studies suggest that both males and females are susceptible to sleep disruption using the ELSD paradigm, however our results suggest that females may be more resilient to developing lasting behavioral impairments (both social and sleep) as a result of this early life sleep disruption than males.

As research in adults with ASD suggest that the presence and/or degree of adult sleep disturbance correlates with the severity of autism symptomology[35, 37–40] the milder social phenotype associated with female ELSD voles compared with males [REF Jones 2019] is perhaps consistent with the lack of sleep disturbances in adult ELSD females.

### Potential limitations, and implications

There are several limitations of the current work to be considered when interpreting the data presented here. Importantly, sleep was recorded from tethered EEG and EMG electrodes that require single housing in order to avoid damaging cables. Prairie voles are highly social rodents and it is possible that this period of single housing influenced their natural sleep behavior. We also did not conduct social or cognitive testing in these experiments, future work will examine correlations between social bonding and cognitive flexibility and REM sleep measurements. Finally, our studies conducted in Experiment 1 were not powered to detect sex differences during ELSD. It is important to know if sleep is affected equally in males and females during the ELSD paradigm. Although the sample size was small for each sex, we did not observe any notable sex differences to support expanding the animal usage for this experiment. Future work using wireless telemetry devices to acquire EEG and EMG signals during sleep will be considered to more thoroughly track sleep ontogeny from the beginning of the ELSD period into young adulthood in both male and female prairie voles thereby expanding on the developmental time points chosen here.

## Conclusion

We find that early life sleep disruption for 1 week during postnatal development in the highly social prairie vole rodent chronically reduces REM sleep duration and has long lasting effects on the later development of REM sleep in adult males. Combined with our previously published research describing long term effects on male social bonding behavior after early life sleep disruption, these results mirror many of the behavioral and sex specific hallmarks of autism spectrum disorder.

## Abbreviations

REM: Rapid Eye Movement
ELSD: Early Life Sleep Disruption
EEG: Electroencephalography
EMG: Electromyography
NREM: Non Rapid Eye Movement
VA: Veterans Affairs
ANOVA: Analysis of Variance
SEM: Standard Error of the Mean
ASD: Autism Spectrum Disorder

## Acknowledgments

The data in this work was supported with resources and the use of facilities at the VA Portland Health Care System, National Science Foundation NCS Foundations Award # 1926818 to H.C. and M.M.L., NIH NHLBI T32 HL083808-10 to C.E.T., NIH NIDA T32 DA007262 to R.J.O, NIH NIA T32 AG055378 to N.E.M., and Portland VA Research Foundation to M.M.L. The contents do not represent the views of the US Department of Veterans Affairs or the United States Government.

