## Supplemental Figures for "Early life sleep disruption has long lasting, sex specific effects on later development of sleep in prairie voles"

**SUPPLEMENTAL RESULTS**

|  | **Day 1** | **Day 2** | **Day 3** | **Day 4** | **Day 5** | **Day 6** |
| --- | --- | --- | --- | --- | --- | --- |
| Male Control | | | | | | |
| **Mean** | 14.92% | 10.32% | 13.12% | 8.97% | 11.21% | 10.56% |
| *SD* | *2.15%* | *0.61%* | *6.47%* | *5.70%* | *6.46%* | *6.75%* |
| n | 2 | 2 | 2 | 2 | 2 | 2 |
| Male ELSD | | | | | | |
| **Mean** | 4.84% | 7.53% | 5.99% | 6.43% | 7.52% | 8.03% |
| *SD* | *2.19%* | *0.99%* | *1.91%* | *1.08%* | *1.20%* | *1.27%* |
| n | 3 | 3 | 3 | 3 | 3 | 3 |

Table S1: Experiment 1 REM sleep % in males during ELSD or Control conditions

|  | **Day 1** | **Day 2** | **Day 3** | **Day 4** | **Day 5** | **Day 6** |
| --- | --- | --- | --- | --- | --- | --- |
| Female Control | | | | | | |
| **Mean** | 16.29% | 12.34% | 13.10% | 11.49% | 9.32% | 10.79% |
| *SD* | *4.65%* | *2.47%* | *3.61%* | *1.02%* | *2.28%* | *4.10%* |
| n | 3 | 3 | 3 | 3 | 3 | 3 |
| Female ELSD | | | | | | |
| **Mean** | 8.56% | 7.03% | 6.20% | 5.89% | 7.76% | 4.64% |
| *SD* | *2.23%* | *1.81%* | *1.13%* | *1.06%* | *0.18%* | *3.86%* |
| n | 2 | 2 | 2 | 2 | 2 | 2 |

Table S2: Experiment 1 REM sleep % in females during ELSD or Control conditions

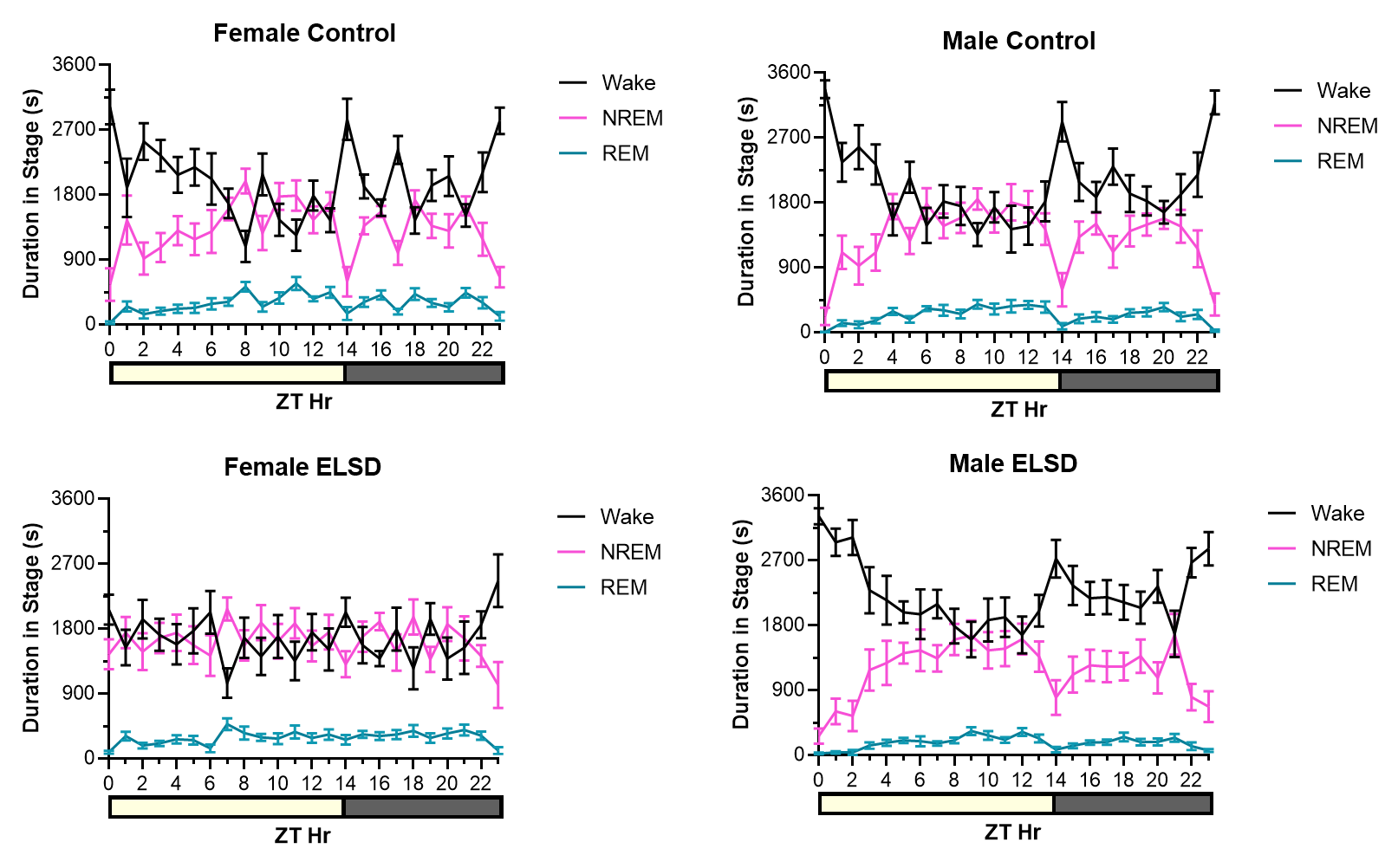

**Fig S1. Adult hourly stage** **duration** **for Control (top panel) and ELSD voles (bottom panel) by sex (females on left, males on right).** Duration in each stage was analyzed by separate repeated measures ANOVA (between subject factors = sleep group, sex; within subjects factor = hour) and there were significant sex x hour interactions for each stage but no within subjects interactions with early life sleep group. Yellow portion of bar represents time in light portion of the light cycle and grey represents dark portion of the light cycle.
